# Predicting the Spatial Dynamics of *Wolbachia* Infections in *Aedes Aegypti* Arbovirus Vector Populations in Heterogeneous Landscapes

**DOI:** 10.1101/458794

**Authors:** Penelope A. Hancock, Scott A. Ritchie, Constantianus J. M. Koenraadt, Thomas W. Scott, Ary A. Hoffmann, H. Charles J. Godfray

## Abstract

1. A promising strategy for reducing the transmission of dengue and other arboviral human diseases by *Aedes aegypti* mosquito vector populations involves field introductions of the endosymbiotic bacteria *Wolbachia*. *Wolbachia* infections inhibit viral transmission by the mosquito, and can spread between mosquito hosts to reach high frequencies in the vector population. *Wolbachia* spreads by maternal transmission, and spread dynamics can be variable and highly dependent on natural mosquito population dynamics, population structure and fitness components.
2. We develop a mathematical model of an *Ae. aegypti* metapopulation that incorporates empirically validated relationships describing density-dependent mosquito fitness components. We assume that density dependence relationships differ across subpopulations, and construct heterogeneous landscapes for which model-predicted patterns of variation in mosquito abundance and demography approximate those observed in field populations. We then simulate *Wolbachia* release strategies similar to that used in field trials.
3. We show that our model can produce rates of spatial spread of *Wolbachia* similar to those observed following field releases.
4. We then investigate how different types of spatio-temporal variation in mosquito habitat, as well as different fitness costs incurred by *Wolbachia* on the mosquito host, influence predicted spread rates. We find that fitness costs reduce spread rates more strongly when the habitat landscape varies temporally due to stochastic and seasonal processes.
5. Our empirically based modelling approach represents effects of environmental heterogeneity on the spatial spread of *Wolbachia.* The models can assist in interpreting observed spread patterns following field releases and in designing suitable release strategies for targeting spatially heterogeneous vector populations.

## INTRODUCTION

*Aedes aegypti* mosquitoes transmit several arboviruses, including dengue which infects an estimated 390 million people per year (Bhatt *et al.* 2013), as well as Zika and chikungunya which have expanded geographically on a global scale in recent years (Nunes *et al.* 2015; Faria *et al.* 2017). Vaccines against these viruses are not yet widely available (Saez-Llorens *et al.* 2017) and disease control strategies rely on interventions that target the mosquito vector. A promising new control strategy involves the field releases of the endosymbiotic bacteria *Wolbachia* into *Ae. aegypti* populations*. Wolbachia* infections in *Ae. aegypti* inhibit virus development and their transmission by the vector (Moreira *et al.* 2009; Walker *et al.* 2011). *Wolbachia* is maternally transmitted in the mosquito host, and utilizes a mechanism known as cytoplasmic incompatibility (CI), which drives the invasion of the bacteria throughout the population. CI works by preventing the development of offspring from matings between *Wolbachia-* infected males and uninfected females (Caspari & Watson 1959), conferring a strong relative fitness advantage to infected females that are able to mate with both infected and uninfected males. Field releases conducted in northeast Australia in 2011 achieved the first successful invasion and stable establishment of a *Wolbachia* strain (*w*Mel) in wild *Ae. aegypti* populations (Hoffmann *et al.* 2011). Current field trials are evaluating the public health impact, as measured by reductions in human disease incidence, of *Wolbachia* infected *Ae. aegypti* (O’’Neill *et al.* 2018).

Given the growing need and potential for applying *Wolbachia* to mosquito vector control and disease prevention, it is necessary to understand the dynamics of *Wolbachia* spread in field *Ae. aegypti* populations. The spatial spread of *Wolbachia* is expected to depend on the natural heterogeneity in mosquito abundance and demography that arises from changing ecological and environmental conditions (Hancock *et al.* 2016a; Hancock *et al.* 2016b). Theoretical studies demonstrate that spatial heterogeneity in host abundance is critical to *Wolbachia* spread, slowing and sometimes stopping the invasion (Barton 1979; Hancock & Godfray 2012). Demographic variation amongst individual hosts, and how this is spatially structured, also has important consequences for spread because the rate of maternal transmission of *Wolbachia* depends directly on its hosts’ fitness (Hancock *et al.* 2016b).

In mosquito populations, important demographic traits such as female fecundity and larval development rates can vary dramatically depending on the levels of density-dependent competition for limited food resources that are experienced during larval development (Barrera *et al.* 2006; Muriu *et al.* 2013; Hancock *et al.* 2016a). Parameters describing female fecundity and larval development rates are fundamental to models of *Wolbachia* dynamics in mosquito populations, because the bacteria is only transmitted maternally (Turelli 1994). Incorporation of density-dependent demographic variation in these parameters into models of the spatial dynamics of *Wolbachia* has, however, been hindered by the lack of an empirical understanding of the form of density dependence (Legros *et al.* 2009). Models of the spatial spread of *Wolbachia* in mosquito populations developed thus far have typically either assumed constant demographic rates across space and time (Turelli & Hoffmann 1991; Schmidt *et al.* 2017; Turelli & Barton 2017), or restricted the effects of density-dependence to impacts on larval survival (Hancock & Godfray 2012).

Here we develop a spatially explicit model of *Wolbachia-Ae. aegypti* dynamics that aims to incorporate realistic patterns of demographic variation in the mosquito population. *Ae. aegypti* is a highly domestic species that is found in or near areas with human habitation and has low rates of dispersal (Harrington *et al.* 2005), which makes population size estimation easier than for other mosquito species (Ritchie *et al.* 2013a). Studies from multiple geographic regions show characteristic strong spatial overdispersion in population size (Getis *et al.* 2003; Focks & Alexander 2006; Koenraadt *et al.* 2008; Jeffery *et al.* 2009; Aldstadt *et al.* 2011; LaCon *et al.* 2014), which may arise from the reliance of *Ae. aegypti* larvae on developing in water filled artificial containers close to human dwellings (Southwood *et al.* 1972; Aldstadt *et al.* 2011; Williams *et al.* 2013). Many of these containers produce low numbers of pupae (and adults), which is thought to be due limited larval food resources (Arrivillaga & Barrera 2004; Focks & Alexander 2006). Adult body size in field populations is typically smaller than those produced from larvae that are fed *ad libitum* in the laboratory (Nasci 1986; Barrera *et al.* 2006), with more crowded field containers typically producing smaller adults (Schneider *et al.* 2004). This indicates that larval development in field *Ae. aegypti* populations is affected by food limitation arising from density-dependent larval competition, which adversely affects juvenile and adult fitness components (Hancock *et al.* 2016a). Thus, density dependent effects on mosquito production and fitness could be an impediment for *Wolbachia* to establish and spread, especially as releases of additional mosquitoes will lead to increased crowding in containers.

*Wolbachia* spread may also be impacted by spatio-temporal variation in mosquito habitat (Schmidt *et al.* 2017). Field studies have shown that over time it is not always the same houses that are responsible for the production of high numbers of *Ae. aegypti* (Jeffery *et al.* 2009; LaCon *et al.* 2014), suggesting that the quality of mosquito habitat within a given house may be temporally variable. In some environments, including northeast Australia, *Ae. aegypti* abundance is seasonal and higher in the wet season (November-April) than in the dry season (May-October) (Ritchie *et al.* 2013b). In some other environments, however, *Ae. aegypti* abundance does not show seasonal variation (Scott *et al.* 2000b; Koenraadt *et al.* 2008).

We develop a metapopulation model that describes the spatially heterogeneous demography of *Ae. aegypti* using empirically validated relationships that link density-dependent demographic traits to mosquito abundance. Incorporating an empirically realistic form of density-dependence significantly reduces parameter uncertainty compared to other models of *Ae. aegypti* population dynamics. We show that our model can produce spatially aggregated patterns of demographic variation similar to those observed in field populations, and that it predicts rates of spatial spread of *Wolbachia* close to those observed following field releases. We then explore the effects on the rate of spread of different types of spatio-temporal variation in the mosquito habitat landscape, including stochastic and seasonal variation, and of different fitness costs incurred by *Wolbachia* on the mosquito host.

## METHODS

We develop an age-structured model of an *Ae. aegypti* metapopulation that assumes that variation amongst individuals in two demographic traits, larval development times and per-capita female fecundity, is influenced by varying levels of larval density-dependent competition for limited food resources (Hancock *et al.* 2016b). The level of density-dependent competition within each of the subpopulations that make up the metapopulation is assumed to depend on the local larval density as well as the quantity of larval food resources available to the subpopulation (Fig 1A). We conceptualize each subpopulation of the metapopulation as a “house” and define the larval habitat quality in each house as high quality, low quality or empty (no larval habitat present). High quality habitats contain a greater larval food resource quantity than low quality habitats.

**Figure 1.**
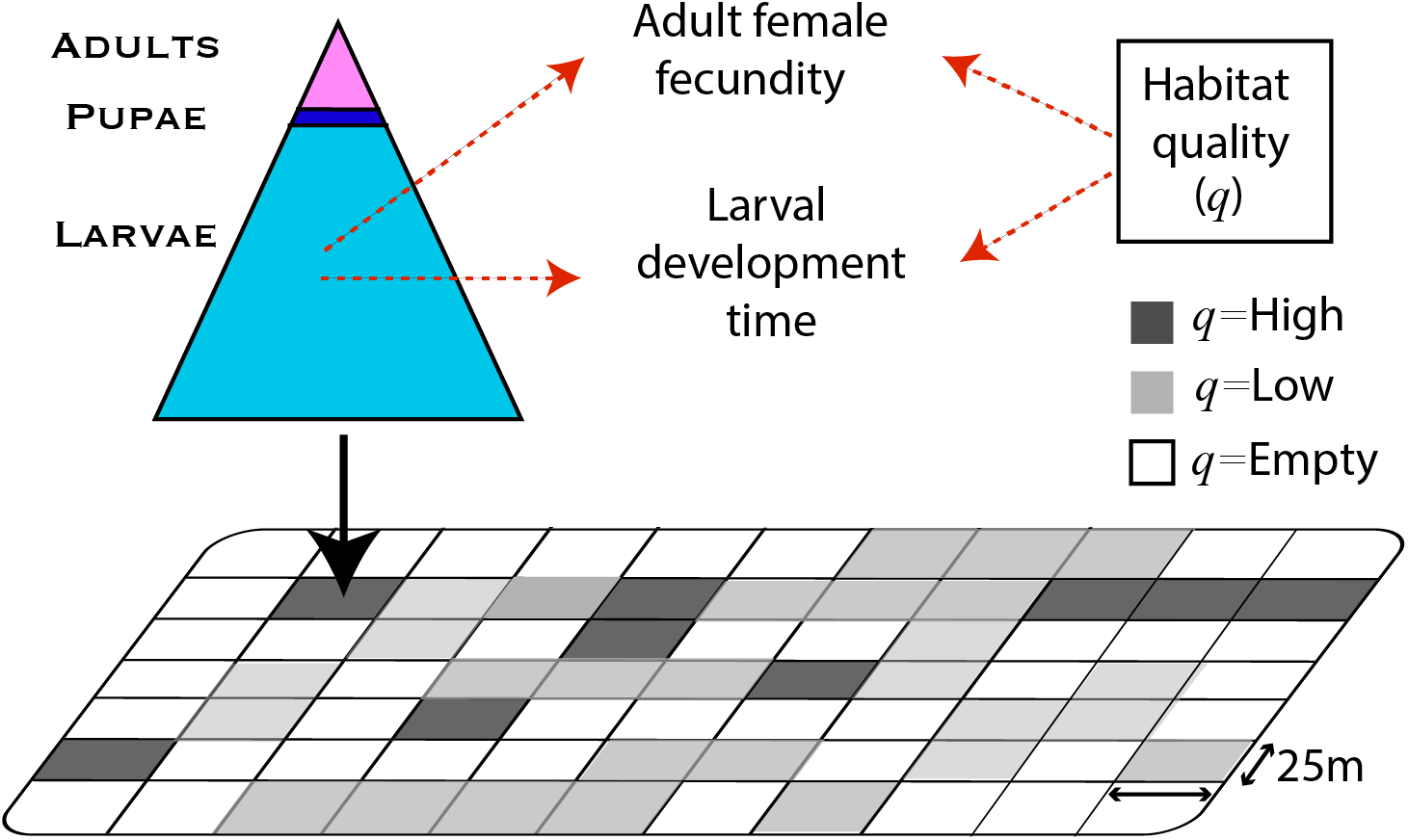
Diagram of the metapopulation model showing interactions between mosquito demographic traits, larval density and habitat quality across subpopulations.

In order to estimate the functional form of density-dependent demographic variation, we utilize a recently developed methodology that is based on frequent observations of the abundance of the juvenile life stages of *Ae. aegypti* in two experimental field-caged populations over a period of approximately six months (see below and Hancock et al. (2016b)). One field-caged population received a higher quantify of larval food resources per week and the other a lower quantity (see below). Mathematical functions describing the density-dependent co-variation in per-capita female fecundity and larval development times occurring in each population are then parameterized from the observed variation in juvenile abundance (Hancock *et al.* 2016b). We use these two sets of relationships to represent the form of density dependence occurring in the high and low quality habitats in our metapopulation model. The other model parameters, including those describing density-independent mosquito demographic traits and dispersal, are assumed to be constant and are estimated using published data.

We describe below the details of our methodology for estimating the density-dependent and density-independent model parameters. Parameters values are provided in Tables 1 & 2 and the mathematical formulation of the model is given in Section S4. We then present approaches for comparing patterns of demographic variation in our simulated metapopulation to those observed in field *Ae. aegypti* populations. Finally we describe our method for comparing predicted rates of spatial spread of *w*Mel *Wolbachia* to those observed in the field.

**Table 1.** Values of model parameters describing mosquito demographic traits and dispersal.

| Symbol | Definition | Value |  | Source |
| --- | --- | --- | --- | --- |
| $T_P$ | Duration of the pupal stage | 2 days | | (Hancock et al. 2016a) |
| $T_G$ | Minimum time for females to become gravid following emergence | 4 days | | (Hancock et al. 2016a) |
| $T_H$ | Time between oviposition and hatching of eggs | 6 days | | (Hancock et al. 2016a) |
| $\omega$ | The proportion of uninfected offspring produced by *Wolbachia*-infected females | 0.01 | | (Walker et al. 2011) |
| $s_h$ | The proportion of unviable offspring from an incompatible mating | 0.99 | | (Walker et al. 2011) |
| $s$ | The reduction in average adult lifespan caused by *Wolbachia* carriage | varies | | |
| $\alpha, \beta, \gamma$ | Parameters of functions describing mean larval development time | *Low quality habitats* | *High quality habitats* | This study |
|  |  | 19.6, 2.74, 0.43 | 1.2, 0.154, 0.68 |  |
| $\nu, \eta, \psi$ | Parameters of functions describing the standard deviation of larval development time | -0.97, 1.0, 0.32 | 12, 0.24, 0.44 | This study |
| $a, b$ | Parameters of functions describing per-capita female fecundity | 15.4, -2.03 | 28.5, -3.25 | This study |
| $\lambda_{\min}, \mu_{\max}, \sigma_{\max}$ | Lower limit on per-capita female fecundity, and upper limits on larval development time means and standard deviations | 0.5, 60 days, 40 days | | This study |
| $\mu_L^c$ | Daily larval mortality | 0.05 | | (Hancock *et al.* 2016a) |
| $\mu_A^c$ | Daily pupal and adult mortality | Field-cage | Field release simulations | (Muir & Kay 1998; Walker *et al.* 2011) |
|  |  | 0.03 | 0.1 |  |
| $p_d$ | Daily probability that adults move to adjacent house | 0.56 | | (Harrington *et al.* 2005) |

**Table 2.** Values of model parameters describing spatial structure and the sizes of mosquito releases.

| Symbol | Definition | Value |
| --- | --- | --- |
| $x_h$ | Side length of square grid units (houses) | 25 m |
| $X_A$ | Side length of square spatial region containing the target mosquito vector population | 2.5 km |
| $h_H, h_L, h_E$ | Proportions of houses with high, low and empty habitat quality | 0.1, 0.45, 0.45 |
| $n_r$ | Number of releases | 15 |
| $t_r$ | Duration of release period | 15 weeks |
| $s_r$ | Number of mosquitoes released per release house per release* | 5* |
| $X_r$ | Size of the release zone* | 725 m<sup>2</sup> |
| $f_r$ | Wolbachia frequency in the release zone at the time of the final release* | 0.85 |

\*Used for model simulations where the fitness cost $s=0$ and the habitat quality landscape was static.

### Estimating the form of density-dependent demographic variation

We estimated the forms of density-dependent variation in mosquito demographic traits occurring in two independent experimental populations of *Ae. aegypti*. Each population was initiated with a cohort of 100 first instar larvae that were the progeny of wild caught *Ae. aegypti* collected in Cairns, northeast Australia. The populations were housed in field-cages designed to simulate the natural habitat of *Ae. aegypti* and the climatic conditions of northeast Australia (Darbro *et al.* 2012) and maintained as described in Hancock *et al.* (2016a; 2016b) (see Section S1 of the Supplementary Material). Each population received a different larval food regime. One population, which we refer to as the HF (high food) Population, received a fixed quantity of food (0.32g ground lucerne) three times per week, and the other, referred to as the LF (low food) Population, received this same food quantity only once per week.

We monitored the abundance of larvae and pupae over approximately six months (194 days for the HF Population and 187 days for the LF Population). All newly eclosed pupae were counted daily and all larvae (categorized as first, second, third and fourth instar) were counted three times per week. We used Bayesian Markov Chain Monte Carlo (MCMC) methods, described in Hancock *et al.* (2016b), to estimate the parameters of mathematical functions describing the form of larval density-dependent variation in larval development times and per-capita female fecundity for each population. These methods calculate the likelihood of our observed abundances of the juvenile mosquito life stages over time. Details of this methodology are provided in Section S2. For both populations, the fitted models accurately describe the major features of the temporal variation in juvenile mosquito abundances in the field-caged populations (Fig S1). For the HF Population, these model fits have been reported previously (Hancock *et al.* 2016b), while for the LF Population our results are reported here for the first time. We note that incorporating density-dependent variation in additional demographic traits, such as the daily larval mortality rate, may have produced a closer fit to the field cage observations.

### Estimating density-independent demographic traits

Values of parameters describing density-independent demographic rates (assumed constant) were estimated from published studies (Table 1). We estimated the daily probability, *p_d_*, that adults move to an adjacent house (within the Moore neighbourhood), using published data from mark-release-recapture (MRR) experiments performed in Thailand (lines 1 and 2 of Table 2 of Harrington et al. (2005)) that record numbers of recaptured *Ae. aegypti* mosquitoes at different distances from the release sites. We simulated the MRR protocol described in Harrington et al. assuming a constant daily rate of adult mosquito mortality *μ_A_* (Table 1). We calculated the value of *p_d_* that provided the minimum least squares fit to the observed recapture rates. We obtained *p_d_* = 0.56, which is higher than the estimated daily movement probability of 0.3 obtained by Magori et al. (2009), who also simulated the Harrington et al. MRR data but defined the set of neighbouring houses as a von Neumann neighbourhood. We assumed that larvae experience a constant daily rate of mortality *μ_L_* similar to the value estimated for our field-cage experiments (Hancock *et al.* 2016b), and that pupae experience the same daily mortality rate as adults. Values of the development time lags *T_G_*, *T_H_* and *T_P_* were estimated from our field-cage experiments (Hancock *et al.* 2016b).

### Estimating spatial variation in habitat quality

We denote the proportion of houses in the metapopulation with high habitat quality, low habitat quality and empty as *h_H_*, *h_L_*, and *h_E_*. We assume that the habitat quality type across houses is spatially uncorrelated (Getis *et al.* 2003) and is described by a multinomial distribution Mult(*h_H_*, *h_L_*, *h_E_*). We then compare the abundance of pupae across subpopulations simulated by our model with observations from field surveys conducted by Koenraadt et al. (2008). They recorded the number of pupae found across a set of residential properties distributed over two villages in Kamphaeng Phet province, Thailand, in four surveys conducted over 2 month periods at the end of the dry season (March and April, which is the beginning of the dengue transmission season) and at the end of the wet season (September and October, which is the end of the dengue transmission season) in 2004 and 2005. They made a total of 2123 observations across 604 houses. For each house, pupae in all water-holding containers were counted. Pupal abundances were determined for all potential larval development sites. The observed distribution of pupal numbers across houses across both surveys shows a spatially aggregated pattern that is typical for *Ae. aegypti* field populations (Getis *et al.* 2003; Focks & Alexander 2006; Jeffery *et al.* 2009; LaCon *et al.* 2014).

We ran one-year simulations of our metapopulation model across which the only parameters that varied were the proportions of houses with high quality, low quality and empty habitats (*h_H_*, *h_L_*, and *h_E_*). We calculated the parameters of the multinomial habitat quality distribution for which the simulated distribution of pupal numbers per house best matched that observed (Table 2). Our approach assumed that the habitat quality of each house does not vary over time (see the Discussion). Further details of this methodology are provided in Section S3.

### Comparing the predicted intensity of density dependence with observations from field populations

We assessed how the intensity of larval density-dependent competition occurring in our modelled metapopulation compared to observations from field populations using observed wing lengths of adult females produced from field-collected *Ae. aegypti* pupae. Food limitation during larval development greatly affects overall adult body size, which can be estimated using wing length (Briegel 1990; Barrera *et al.* 2006), so field populations that show smaller adult wing lengths are likely to have experienced more intense larval resource competition (Barrera *et al.* 2006). Temperature conditions can also affect *Ae. aegypti* body size (Scott *et al.* 2000a; Mohammed & Chadee 2011), although in our field-cage populations changes in larval water temperature had less influence on demographic traits than variation in larval density (Hancock *et al.* 2016a) (see the Discussion). We used published observations of the average wing length per household of 2316 female *Ae. aegypti* collected as pupae from 2931 houses in Iquitos, Peru (Schneider *et al.* 2004). The wing lengths of the sampled mosquitoes are highly variable, and median wing length is low relative to that occurring when adult females experience plentiful food conditions during larval development (Fig. 1 of (Schneider *et al.* 2004)). Other observed wing lengths distributions obtained from field-sampled *Ae. aegypti* also show similar characteristics (Nasci 1986; Barrera *et al.* 2006).

In order to compare these wing length observations to the output of our model, we used the relationship between wing length and female fecundity obtained by Briegel (1990). Briegel (1990) recorded adult female wing length, *w*, and the number of eggs oviposited per gonotrophic cycle, *λ*̃_*G*_, for 206 *Ae. Aegypti* females reared in a laboratory under varying larval density conditions and blood-fed on human hosts (Table 3 of Briegel 1990))). He then fitted a linear relationship to these observations:

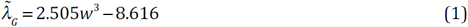

We applied eqn (1) to estimate the distribution of the average *λ*̃_*G*_ per house, *λ_G_*, *_h_*, from the observed distribution of average female wing lengths per house. This procedure assumes that individual wing lengths can be approximated by the house average. We then calculated the distribution of the weekly average per-capita female fecundity for each house, *λ_h_*, using observations from our field-cage experiments (see Methods). This procedure makes two further assumptions: (i) *λ_G_*, *_h_* and *λ_h_* are assumed to be directly proportional; and (ii) the adult female wing lengths in our field-caged populations under conditions of plentiful larval food are equal to the highest value observed by Briegel (1990). Having obtained a distribution of per-capita female fecundity across individuals derived from the field wing length data, we compared it with the distribution of per-capita female fecundity values across all subpopulations simulated by our metapopulation model.

### Comparing the predicted rate of spatial spread of *w*Mel *Wolbachia* with observations from field populations

We compared the rates of spatial spread of *Wolbachia* predicted by our model to those observed following the *w*Mel field release trials conducted at two sites in the city of Cairns, northeast Australia, in 2013 (Schmidt *et al.* 2017). The two sites, Edge Hill and Parramatta Park, are residential areas close to the center of Cairns. The field trials involved releasing mosquitoes infected with *w*Mel into a restricted area, or “release zone”, with the aim of allowing the *Wolbachia* to spread into surrounding areas. Observed rates of spread of *w*Mel beyond the release zone for both sites are reported in Schmidt et al. (2017) (see Methods). We used their Gaussian/logistic model fits to observations of *w*Mel frequencies in pooled samples (Fig. S3 of Schmidt et al. (2017)) to obtain the estimated distance from the edge of the release zone at which the *w*Mel frequency was 0.5 at multiple time points following the releases.

To represent their spatial release strategy, we assume in our model that mosquitoes infected with *w*Mel are liberated within a square-shaped release zone located at the center of a larger area of potential habitat containing 10,000 houses (also square-shaped; 2.5 km^2^). We set parameters specifying the release protocol to values matching those used in the field trials (these included the area of the release zone, the duration and frequency of releases, and the frequency of *w*Mel inside the release zone at the time of the final release; Table 2). We assumed that a fixed number of infected mosquitoes are released within each house inside the release zone once per week. Because released mosquitoes were reared in the laboratory, we also assumed that the per-capita fecundity of the released females is high and equal to that observed when larval food resources are plentiful (Hancock *et al.* 2016b). Our results assume that houses have dimensions of 25m^2^ (Table 2), which is approximately equal to average area of the housing blocks within the two release zones in Cairns, calculated by randomly sampling 100 houses within each release zone and obtaining the dimensions of each housing block using Google Maps.

Our model of the *Wolbachia* dynamics in each subpopulation of the metapopulation is based on that described in Hancock et al. (2016b), and we provide the mathematical details in Section S5. We assume high values of the strength of cytoplasmic incompatibility, *s_h_*, and the rate of maternal transmission, *ω* (Table 1), consistent with observations of the *Wolbachia* strain *w*Mel that has been used in recent field release trials (Hoffmann *et al.* 2011). We begin by assuming that *Wolbachia* infection does not affect the fitness of its mosquito host (Hancock *et al.* 2016b), and then explore the effects of varying fitness costs on spread dynamics.

## RESULTS

### Modelling spatially varying demography in the mosquito population

#### Modelling spatial variation in mosquito abundance

The fitted distribution of the numbers of pupae across houses is similar in both shape and range to that obtained from the field surveys (Fig. 2A). The predicted pupal numbers are highly aggregated across space due to the small proportion of houses with high quality habitat (*h_H_* = 0.1). The fitted proportions of houses with low and empty habitat quality (*h_L_* and *h_E_*) were equal (Fig. 2A). These results demonstrate that the relationships describing density-dependent demography derived from our field-caged population experiments can be applied directly, without rescaling, to approximate patterns of variation in abundance that occur in field *Ae. aegypti* populations.

**Figure 2.**
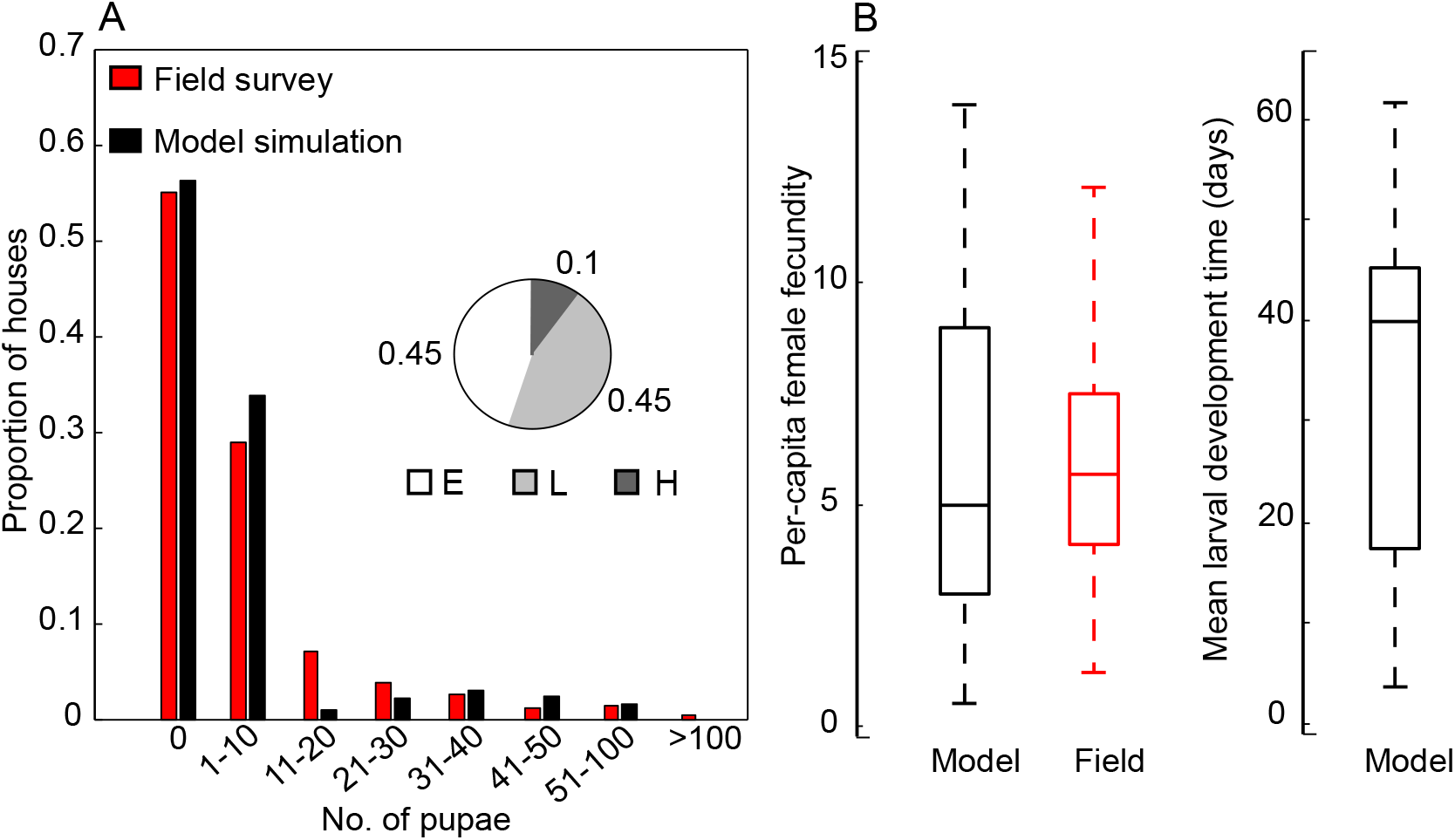
**(A)** The distribution of the number of pupae per house. Red bars show observations from the field surveys and black bars show the simulated distribution given by our metapopulation model. The pie chart shows the fitted proportions of houses with high (H), low (L) and empty (E) habitat quality (*h_H_*, *h_L_*, and *h_E_*, respectively). **(B)** Boxplots showing the distribution of per-capita female fecundity across subpopulations predicted by our model (black, left), the per-capita fecundity distribution estimated from field-caught mosquitoes (red; see text), and the distribution of the mean development times of the larval cohorts predicted by our model (black, right).

#### Modelling the intensity of density-dependent competition

The median of the field-derived per-capita female fecundity distribution is in the lower part of the per-capita fecundity range and is close to that predicted by our model (Fig. 2B). Thus both field observations and model predictions indicate that density-dependent competition is relatively intense. While both the distributions from the field and model show wide variation in per-capita female fecundity across subpopulations, the variance predicted by our model is larger. The larval development times predicted by our model also vary widely across the metapopulation, with the majority of subpopulations having protracted larval development (Fig. 2B).

### Predicting the speed of spatial spread of *Wolbachia*

We simulated *Wolbachia* field releases for 50 landscapes where habitat quality in each house is randomly drawn from a multinomial distribution with parameters *h_H_*, *h_L_*, and *h_E_* (Table 1). The mean of the simulated rates of spread across all landscapes agrees well with the rates observed at Edge Hill and Parramatta Park (Fig. 3A). For each landscape the simulated spread rates beyond the release zone were spatially heterogeneous (Fig. 3A and see Animation S1) and the spatial spread pattern varied across the different landscapes.

**Figure 3.**
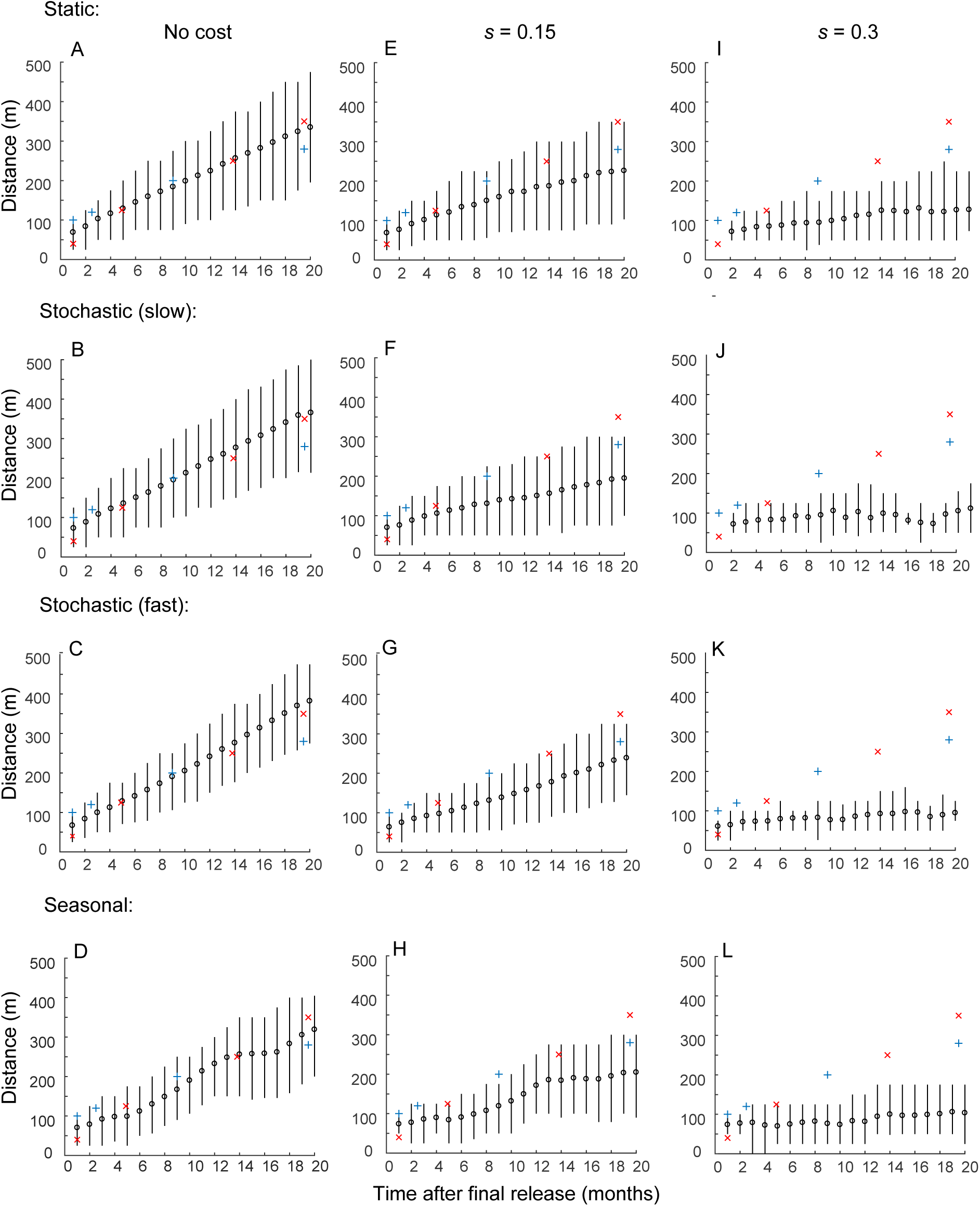
Comparing predicted and observed rates of spatial spread of *w*Mel *Wolbachia*. Black circles show the mean across all simulations of the distance of the 0.5 *w*Mel frequency contour from the edge of the release area and vertical black lines connect the 5% and 95% quantiles. The observed mean position of the 0.5 *w*Mel frequency contour from the edge of the release area following the *w*Mel releases conducted in Cairns is shown for two release areas, Edge Hill (red crosses) and Parramatta Park (blue crosses). Four types of temporal variation in house habitat quality are shown: Static habitat quality (Top row **(A,E,I)**); Slow stochastic temporal variation (*t* ̅ *_q_* =3 months; Second row **(B,F,J)**); Fast stochastic temporal variation (*t*̅*_q_* =1 month; Second row **(C,G,K)**); Seasonal variation (Fourth row **(D,H,L)**). Three values of the fitness cost incurred by *Wolbachia* are shown: No cost (*s*=0; First column **(A-D)**); *s*=0.15 (Second column **(E-H)**); *s*=0.3 (Third column **(I-L)**).

#### Effects of stochastic habitat variation

We first extend our analysis to consider landscapes where habitat quality shows stochastic temporal variation. We defined a fixed daily probability *t_q_* that the habitat quality in each house is reassigned to a value chosen randomly from Mult(*h_H_*, *h_L_*, *h_E_*). We then defined the expected habitat quality turnover period, *t* ̅ *_q_* = 1 / *t_q_*, as the expected time interval between a reassignment of habitat quality for each house, and considered cases where turnover is slow (*t*̅ *_q_* =3 months) and relatively fast (*t*̅ *_q_* =1 month). For each value of *t*̅ _*q*_ we repeated the analysis described in the previous section to obtain predictions of the speed of spatial spread of *Wolbachia* by simulating the release protocol used in the field releases conducted in Edge Hill and Parramatta Park, Cairns (Table 2). We found that the predicted rate of spatial spread of *Wolbachia* for both values of *t_q_* was similar to that obtained when habitat quality was assumed not to vary temporally (Fig. 3B-C).

#### Effects of seasonal habitat variation

We next extended our analysis to explore effects of seasonal variation in *Ae. aegypti* abundance on *Wolbachia* spread. We simulated seasonal variation in *Ae. aegypti* abundance by periodically increasing the proportion of high quality habitat, *h_H_*. This involved randomly selecting a set of houses (of size *n_s_*) with either low or empty habitat quality, and reassigning the habitat quality in these houses to high, once every 365 days. This “wet season” habitat quality landscape remained fixed for 182 days, and then a set of high quality habitat houses, also of size *n_s_*, was selected at random and the habitat quality in these houses was reassigned to either low, or empty, with a probability of 0.5. We set the value of *n_s_* such that the proportion of houses with high quality habitat, *h_H_*, increased to 0.175 at the start of the wet season (and returned to 0.1 at the start of the dry season). This produced a simulated pattern of seasonal variation in adult abundance with wet season peak about 4 times higher than the dry season trough (Fig. S3). Published observations of the seasonal variation in the numbers of *Ae. aegypti* caught in Biogents Sentinal traps (BGS) throughout Cairns show a similar relative amplitude across seasons (Fig. 1 of Ritchie et al. (2013b)). We again simulated the above *Wolbachia* field release protocol, starting the releases 40 days into the wet season, and again found that the predicted rate of spatial spread of *Wolbachia* was similar to that obtained when habitat quality was assumed not to vary temporally (Fig. 3D).

#### Effects of fitness costs

We then investigated how these spread patterns are affected by fitness costs imposed by *Wolbachia* on its mosquito host. Specifically, we assumed that *Wolbachia*-infected individuals experience a reduction in average adult lifespan relative to uninfected individuals by a proportion *s.* Estimates of fitness costs incurred by *w*Mel on *Ae. aegypti* are highly uncertain (Hoffmann *et al.* 2011; Hancock *et al.* 2016b; Schmidt *et al.* 2017) (and see the Discussion) but high costs are unlikely; here we consider values of *s*=0.15 and *s*=0.3. For all four types of temporal variation in mosquito habitat quality considered above (including static landscapes with no temporal variation), we simulated the *Wolbachia* release protocol previously described, but augmented the total release size so that the *Wolbachia* frequency reached the required value at the time of the final release (Table 2).

For environments where the habitat quality landscape is static, either fitness cost resulted in slower rates of *Wolbachia* spread that show less agreement with the rates observed in the field populations (Figs. 3E & 3I). When *s*=0.15 the observed spread rates are above the mean of the simulated rates, but within the 95% quantiles (Fig. 3E), and when *s*=0.3 the observed rates are higher than the quantiles of the simulated rates for times later than six months following the final release (Fig. 3I).

In the presence of fitness costs, predicted *Wolbachia* spread rates also differ across the different types of spatio-temporal variation in mosquito habitat quality. They are slower in landscapes where house habitat quality undergoes stochastic changes over time compared to landscapes where house habitat quality is static, with differences being larger when the fitness cost is higher (Figs. 3F-G & 3J-K). Seasonal variation in mosquito habitat quality also results in slower predicted spread rates, with the difference again being strongest for higher fitness costs (Figs. 3H & 3L). When the *Wolbachia* imposes fitness costs, these types of temporal variation in habitat quality result in slower rates of spatial spread because a reduction in a house’s habitat quality causes a localized increase in mosquito mortality. If this occurs before the *Wolbachia* has spread to fixation, the frequency dependent advantage that the *Wolbachia* has gained by invading and spreading in areas with good habitat quality is lost; thus spread slows down, and may fail if frequencies fall too low. Interestingly, the higher rates of mosquito mortality in temporally variable habitat quality landscapes lead to reductions in the intensity of density-dependent larval competition, which can lead to faster *Wolbachia* spread rates (Hancock *et al.* 2016b). However in the presence of fitness costs this effect did not compensate for the inhibitory effects on rates of spread of temporal changes in house habitat quality.

## DISCUSSION

Our metapopulation model describes the dynamics of *Ae. aegypti* populations with spatially heterogeneous abundance and demography, incorporating empirically derived relationships that model density-dependent demographic traits. We demonstrated that the model could approximate the patterns of demographic variation that occur in field populations and predict rates of spatial spread of *Wolbachia* close to those observed following field releases. We discuss below how our representation of spatially structured population dynamics is important to predicting *Wolbachia* spread, and also identify limitations of our model in describing the dynamics of natural populations.

Previous studies have demonstrated several mechanisms through which the dynamics of *Wolbachia* spread are affected by spatial heterogeneity in host abundance (Barton 1979; Hancock & Godfray 2012), and demographic heterogeneity across individuals (Hancock *et al.* 2016a; Hancock *et al.* 2016b). Our modelling approach empirically relates variation in mosquito abundance to density-dependent variation in two demographic traits, namely larval development times and per-capita female fecundity, which are major determinants of the rate of *Wolbachia* transmission through successive (overlapping) mosquito generations. By allowing density dependence relationships to vary across subpopulations, our model is able to represent realistically two important demographic features of *Ae. aegypti* field populations: (i) highly variable, spatially aggregated patterns of abundance; and (ii) a relatively intense level of density-dependent larval resource competition on average across the metapopulation.

However, other aspects of demographic and environmental variation that were not included in our model may also cause variable rates of *Wolbachia* spread. Mosquito mortality and dispersal rates also affect *Wolbachia* spread dynamics (Schofield 2002; Hancock *et al.* 2016b), but these traits are poorly quantified and likely to vary with changing environmental conditions (Harrington *et al.* 2005; Maciel-De-Freitas *et al.* 2007; Hoffmann 2014). Several demographic traits vary due to changes in weather and climate (Scott *et al.* 2000a; Mohammed & Chadee 2011), and predation (Bowatte *et al.* 2013). Moreover, density dependence may impact other demographic traits such as stage-specific daily survival rates (Hancock *et al.* 2016a), and mating success (Segoli *et al.* 2014). Density-dependent dynamics may be further complicated by preferential oviposition behaviour by adult females, whereby females are more likely to oviposit in containers that are already inhabited by conspecific larvae (Wong *et al.* 2011). Thus our model cannot fully describe the demographic variability in a particular metapopulation. By specifying realistic forms of density dependence, however, our model offers a useful tool for exploring dynamic behaviour across different assumptions about uncertain aspects of mosquito-*Wolbachia* demography and different types of environmental variation.

A current challenge in forecasting *Wolbachia* dynamics is the difficulty of estimating fitness costs in the field (Hancock *et al.* 2016b), which is exacerbated by the very high levels of demographic variation caused by environmental and genetic effects (Hoffmann *et al.* 2011; Walker *et al.* 2011; Hoffmann 2014; Hancock *et al.* 2016b; Ross *et al.* 2016; Schmidt *et al.* 2017). This is problematic because theory suggests that even small fitness costs strongly affect the spatial spread of *Wolbachia* (Turelli & Hoffmann 1991; Hancock & Godfray 2012). For example, the classical reaction-diffusion models of Barton and Turelli (Barton 1979; Turelli & Hoffmann 1991) predict that a fitness cost of 0.25 halves the equilibrium wave speed of *Wolbachia*. These reaction-diffusion approaches assume spatial and temporal constancy of mosquito abundance and demography (Turelli & Barton 2017). Under plausible sets of demographic and environmental parameters (Tables 1 & 2), and across a range of models of spatio-temporal habitat variation, our model accurately described observed spread rates without assuming that *Wolbachia* incurs fitness costs. It is important to note that, while *Ae. aegypti* abundance varies seasonally in northeast Australia, in many dengue-endemic regions (such as parts of Thailand and Peru) abundance does not show a strong seasonal signal (Morrison *et al.* 2004; Koenraadt *et al.* 2008). Interestingly, however, for cases where the fitness cost incurred by *Wolbachia* was low, our predicted spread rates were relatively robust to the different forms of seasonal and stochastic habitat variation that we considered. While our results predict that fitness costs incurred by *w*Mel are usually small, the uncertainty surrounding the demographic and environmental parameters and processes that we aim to model prevents us from making accurate inferences about the effects of *Wolbachia* on fitness experienced in field *Ae. aegypti* populations.

In summary, we present a mathematical model of the spatial dynamics of *Wolbachia* in *Ae. aegypti* populations that incorporates realistic features of mosquito demographic variation that impact *Wolbachia* spread. Our approach facilitates further exploration of the effects of demographic and environmental heterogeneity on the spread of *Wolbachia* following field releases. Our model can also be applied to explore the dynamics of other mosquito control interventions, such as the release of *Wolbachia-*infected male mosquitoes in order to suppress vector abundance, or the release of mosquitoes carrying gene drive constructs that aim to suppress abundance or lower vectorial capacity (Flores & O’Neill 2018). By developing our understanding of the ecological dynamics of *Ae. aegypti* arbovirus vector populations, and their interactions with vector control interventions, our models can contribute to interpreting observed impacts in field populations and allowing effects of environmental heterogeneity to be considered in designing intervention strategies.

## AUTHOR CONTRIBUTIONS

PAH, HCG and SAR conceived the ideas; PAH designed the methodology; PAH collected the data; PAH analysed the data; PAH led the writing of the manuscript. All authors contributed critically to the drafts and gave final approval for publication.

## ACKNOWLEDGEMENTS

This research was supported by a Marie Curie International Outgoing Fellowship within the 7th European Community Framework Programme (Grant no. 326551-WOLBACHIA-MOD).

## DATA ACCESSIBILITY

Mosquito age-class abundances for LF Population are available on Figshare. C++ Computer Code for implementing the metapopulation model is available on Figshare.

